# Scene context impairs perception of semantically congruent objects

**DOI:** 10.1101/2020.09.30.320168

**Authors:** Eelke Spaak, Marius V. Peelen, Floris P. de Lange

**Affiliations:** Donders Institute for Brain, Cognition and Behaviour, Radboud University, Nijmegen, The Netherlands

## Abstract

Visual scene context is well-known to facilitate the recognition of scene-congruent objects. Interestingly, however, according to the influential theory of predictive coding, scene congruency should lead to reduced (rather than enhanced) processing of congruent objects, compared to incongruent ones, since congruent objects elicit reduced prediction error responses. We tested this counterintuitive hypothesis in two online behavioural experiments with human participants (N = 300). We found clear evidence for impaired perception of congruent objects, both in a change detection task measuring response times as well as in a bias-free object discrimination task measuring accuracy. Congruency costs were related to independent subjective congruency ratings. Finally, we show that the reported effects cannot be explained by low-level stimulus confounds, response biases, or top-down strategy. These results provide convincing evidence for perceptual congruency costs during scene viewing, in line with predictive coding theory.

**Statement of Relevance:** The theory of the ‘Bayesian brain’, the idea that our brain is a hypothesis-testing machine, has become very influential over the past decades. A particularly influential formulation is the theory of predictive coding. This theory entails that stimuli that are expected, for instance because of the context in which they appear, generate a weaker neural response than unexpected stimuli. Scene context correctly ‘predicts’ congruent scene elements, which should result in lower prediction error. Our study tests this important, counterintuitive, and hitherto not fully tested, hypothesis. We find clear evidence in favour of it, and demonstrate that these ‘congruency costs’ are indeed evident in perception, and not limited to one particular task setting or stimulus set. Since perception in the real world is never of isolated objects, but always of entire scenes, these findings are important not just for the Bayesian brain hypothesis, but for our understanding of real-world visual perception in general.

## Introduction

Objects are typically encountered in particular contexts; e.g. a hair dryer is more commonly encountered in a barbershop than in a greengrocer’s. Semantic associations between real-world scene context and objects within such scenes are well-known to facilitate perception in many circumstances: objects are located and identified more rapidly and accurately within semantically congruent contexts than in incongruent ones (Bar, 2004; Biederman, 1972; Davenport & Potter, 2004; Kaiser et al., 2019; Oliva & Torralba, 2007). These congruency benefits are elegantly explained from the perspective of *predictive coding*, the idea that the brain is a hypothesis-testing machine (Clark, 2013; de Lange et al., 2018; Friston, 2005; Rao & Ballard, 1999): the gist of a (natural) scene induces a prior expectation over particular objects common to such a scene, and stimuli that are likely under that prior are easily integrated with it in order to arrive at a coherent representation.

Such an account explains benefits in cases where the scene-induced expectation is relevant to the task at hand: observers that are asked to locate a computer mouse in a scene of a desk will naturally look for it next to the keyboard, and will thus more quickly find it than if it were presented in a scene of a kitchen countertop. Similarly, in a brief or degraded presentation of such scenes, an oval blob next to a keyboard-shaped blob will more readily be identified as a computer mouse than such a blob in incongruent surroundings. However, according to predictive coding, it is precisely *incongruent* objects that warrant closest inspection, not congruent ones (specifically, *high-precision* incongruent objects warrant closest inspection). The inferred identity of a congruent object is easily integrated with the prior induced by the scene gist, whereas the inferred identity of an incongruent object elicits a larger prediction error. To ‘resolve’ this error (or, equivalently: to leverage the likely high information content afforded by the source of this error signal), incongruent objects should be associated with more extended processing, before integration with the scene-induced prior is possible. It has been demonstrated that the amount of processing influences the level of subjective awareness (Anzulewicz et al., 2015; Windey et al., 2013). Given this assumption and the above reasoning, we hypothesized that, in crowded natural scenes, with a clear gist-induced prior that is not directly relevant for the current behavioural goals, and with multiple objects to be explored (‘sampled’), objects in congruent surroundings should be sampled less and therefore perceived less strongly, less saliently, than those in incongruent surroundings.

Researchers studying change detection have reported such context congruency costs: observers are slower to detect changes in objects when these objects are embedded within congruent contexts, compared to incongruent ones (Hollingworth & Henderson, 2000; LaPointe et al., 2013; Mack et al., 2017). Additionally, it has been reported that, during free viewing, observers tend to fixate earlier on incongruent than on congruent objects (Bonitz & Gordon, 2008; Loftus & Mackworth, 1978; Underwood et al., 2007), and other indices of attentional allocation point in the same direction (Gordon, 2004).

However, this reported evidence for congruency costs (or, equivalently: incongruency benefits) is not clear-cut. First, several studies have failed to replicate the earlier fixation latencies for incongruent objects (De Graef et al., 1990; Henderson et al., 1999), while yet others have reported the effect only for visually non-salient objects (Underwood & Foulsham, 2006). In general, low-level visual saliency has been described as a potentially confounding factor in the research on attentional attraction by semantic incongruence (Underwood & Foulsham, 2006; Võ & Henderson, 2009). A second issue that has received less attention (though see (Hollingworth & Henderson, 2000)), but may be equally grave, is that congruency costs might reflect a *strategic* effect: if an incongruent object is present, in many cases it will be task-relevant (e.g., the changing object in change detection, or something specifically memorable in a memory task), making it beneficial for participants in the experiment to search for incongruent objects in general (leading to an observed congruency cost). Finally, and perhaps most importantly, congruency costs have mainly been demonstrated through latency differences (e.g., change detection latency) rather than through unbiased measures of perception. Response latency is influenced by multiple factors, including decisional and response biases. Previously reported congruency costs may thus reflect such biases, rather than reduced perceptual encoding of congruent objects.

An intriguing possible implication of the influential theory of predictive coding is the existence of congruency costs in purely perceptual (i.e., non-semantic) tasks, yet this hypothesis has not been tested directly. In the present study, we set out to perform this test, while taking care of all three concerns voiced above. Importantly, we examined semantic congruency costs in a discrimination task, probing object-level (i.e., exemplar) perception, free from stimulus, response, semantic, and task-strategic confounds, in addition to a classic change detection setting. In brief, across these two behavioural experiments, with a total of 300 participants, we found that congruency costs (1) are evident in change detection even with a fully balanced stimulus set; (2) generalize to a more directly perceptual identification task; (3) persist even when attending to incongruent objects is strategically disadvantageous; and (4) are explained by the subjective level of object-scene consistency.

## Methods

### Stimuli, task, and experimental design

#### Experiment 1 – change detection

This experiment was a version of the classical change detection “flicker” task (Rensink et al., 1997), and is depicted in Figure 1a. Participants were instructed to detect changes between successive displays of the same scene. Each trial started with a 500 ms empty screen, followed by a fixation button labelled ‘Go’ in the center of the screen that participants had to click in order to initiate visual stimulation. Requiring a mouse click in the center of the screen ensured that participants were always fixating the center at stimulus onset. Stimulation consisted of an object-present scene for 250 ms, followed by a 100 ms blank, followed by an object-absent scene. This sequence was repeated for a maximum of 13 times, or until the participant indicated they had detected the change by pressing the space bar. After the detection response, participants were presented with a grey rectangle of the same dimensions as the scene stimulus, and had to click where they detected the changing object. This ensured task compliance, preventing blind, rapid, space bar pressing.

**Figure 1:**
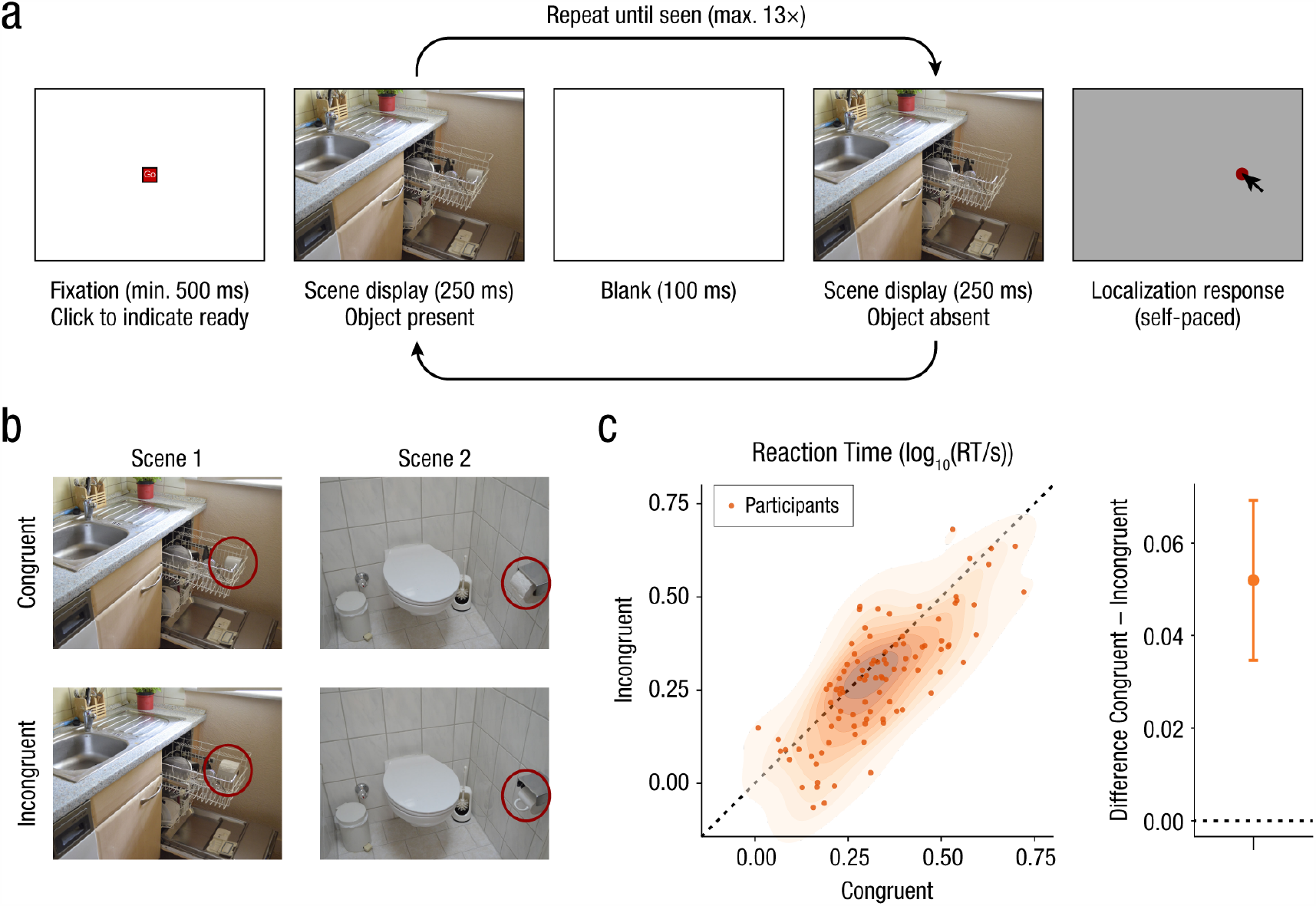
Design and key results for Experiment 1. **(a)** Structure and timeline of a single experimental trial. This was a typical “flicker” change detection task, followed by a localization response. **(b)** Examples of stimuli used in the two conditions (Öhlschläger & Võ, 2017). Note the matched key object (cup/toilet roll in dishwasher versus cup/toilet roll in toilet roll holder). Red outlines indicate the key object (not shown to participants). **(c)** Reaction times across all participants, for Congruent and Incongruent trials. Full distribution in scatterplot, mean ± 95% confidence interval of difference scores on the right. Responses are faster for incongruent than for congruent trials.

Stimuli were taken from a recently published, fully balanced, stimulus database called ‘SCEGRAM’ (Öhlschläger & Võ, 2017). All 62 scenes from this database were used, in the CON and SEM conditions (which we subsequently refer to as the ‘Congruent’ and ‘Incongruent’ conditions), as well as the corresponding object-absent scenes. This database is comprised of pairwise balanced stimuli, matched in lower-level visual features. Each key object occurs in both a congruent context (e.g., a cup in a dishwasher) and an incongruent context (e.g., a cup in a toilet roll holder), and is matched with another key object, similar in shape and orientation, which has the complimentary congruency mapping (e.g., a toilet roll in either a toilet roll holder (congruent), or a dishwasher (incongruent); see Figure 1b).

Each participant completed 62 trials. Half of these were congruent scenes, and half of these incongruent. This mapping was counterbalanced across participants. The main dependent variable (DV) was change detection reaction time, with localization error a secondary DV. Localization error was defined in % of the target object, e.g. a click located 1 cm away from the center of a 1 cm × 1 cm object would be assigned an error of 100%, and any errors < 50% indicate that the observer clicked within the bounds of the target object. Specifically, localization error was defined as:

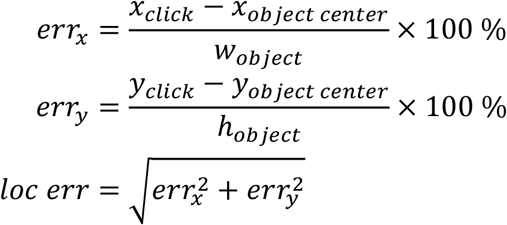

For all analyses of reaction time, we focused only on those change detection responses for which the subsequent localization error was ≤ 100%.

#### Experiment 2 – object identification

For this experiment, instead of having to detect a change, participants were instructed to attentively look at each scene, and afterwards they had to make an identification judgement about an object in that scene. Each trial started with a fixation cross (800 – 1000 ms, randomly drawn from a uniform distribution), followed by a scene display (2.5 s), followed by another fixation cross (500 ms), followed by a two-alternative forced choice (2AFC) response prompt for a maximum of 2.5 s, or until one of the response keys was pressed (Figure 2a). The 2AFC prompt always consisted of the target item that was present in the scene (again taken from the SCEGRAM stimulus database), as well as a lure item, selected through an internet search. The lure was always from the same category as the target, and similar in shape, but a clearly different exemplar.

**Figure 2:**
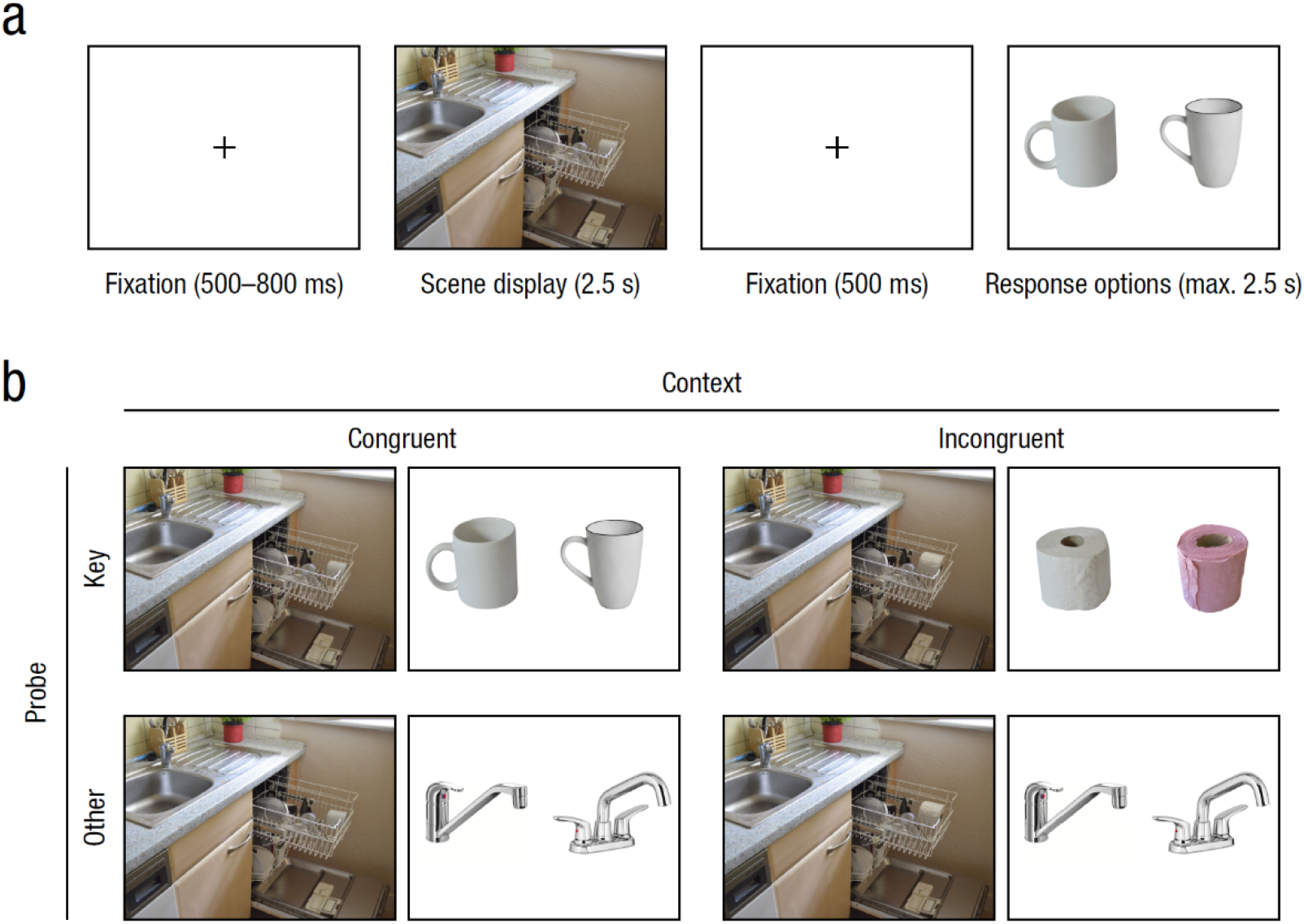
Design for Experiment 2. **(a)** Structure and timeline of a single experimental trial. Participants were presented with a natural (indoor) scene, and were tasked to make a two-alternative forced choice (2AFC) exemplar discrimination afterwards. **(b)** Illustration of the two within-subjects manipulations. Scenes could either be Congruent (left) or Incongruent (right), and the probed objects could either be the Key object (top) or an Other object (bottom).

As in Experiment 1, scenes could occur in a Congruent or Incongruent condition. Whether a scene is Congruent or Incongruent is determined by the nature of what we label the ‘key’ object in the scene (e.g., the cup or toilet roll in Figures 1b and 2). For Experiment 2, we added a ‘Probe-Key’ versus ‘Probe-Other’ factor, governing whether, on a given trial, the participant was probed about this key object, or about another object in the same scene. The probed object in Probe-Other trials was always congruent with the surrounding context (see Figure 2b). The rationale for including Probe-Other trials is twofold. First, this allows us to control for the strategic concern mentioned in the introduction; i.e. if an incongruent object is present, this is no longer necessarily the relevant object (Hollingworth & Henderson, 2000). Second, the presence of an irrelevant congruent/incongruent object might draw attention *away* from the probed target object; this effect should be detectable.

From the participant’s perspective, there is no subjective difference between a Congruent/Probe-Key trial, and a Congruent/Probe-Other trial: on both trial types, there are only congruent items present in the scene, and the participant is later probed about one of them. This distinction nevertheless is important because of the design of the stimulus set. Any given Congruent/Probe-Key stimulus is matched with an Incongruent/Probe-Key stimulus (i.e., the same scene with either a congruent or incongruent item, where these key items are matched for location, shape, and size). Similarly, any given Congruent/Probe-Other stimulus is matched with an Incongruent/Probe-Other stimulus (i.e., again the same scene with a matched congruent/incongruent item present, yet now probed about another, non-manipulated, object, identical across the two levels of congruency). This crossed Probe × Congruency manipulation is very similar to previous work by (Hollingworth & Henderson, 2000).

Stimuli were again counterbalanced across participants, each of whom again completed 62 trials. Trials were again 50% congruent, 50% incongruent. Probe × Congruency together form a 2×2 factorial design, but trial counts per cell were deliberately not fully equalized for all participants. Instead, we introduced a between-subjects factor, *p*(*Probe-Key* = 1 | *Incongruent* = 1), or *p*(*Probe-Key* | *Incongruent*) for short, which governs the proportion of Incongruent trials that are Probe-Key (i.e., given that an incongruent object is present in the scene, *p*(*Probe-Key* | *Incongruent*) determines the probability that this object is the task-relevant one). This factor took on values of 17/33/50/67/83%. For values of the between-subjects factor other than 50% (the fully across-subjects counterbalanced case), a randomly chosen subset (per participant) of Incongruent trials was switched from Probe-Key to Probe-Other, or vice versa. This factor allowed us to quantify the degree to which behavioural costs/benefits were a consequence of task strategy.

The main DV for this experiment was 2AFC accuracy, with reaction time of secondary interest. For reaction time analyses, we focused only on those trials with correct responses.

### Participants, data inclusion, and statistical power

Experiments were performed online, using the Gorilla platform (Anwyl-Irvine et al., 2020), and recruitment was done through the Prolific platform (https://www.prolific.co/). The study was approved by the local ethics committee (CMO Arnhem-Nijmegen, Radboud University Medical Center) under the general ethical approval for online studies for the Donders Centre for Cognitive Neuroimaging.

To increase the signal-to-noise ratio of our data set, we decided *a priori* to remove outliers, which might be especially expected in an online setting. We defined outliers as those participants scoring > 2.5 s.d. away from the mean for either DV. Outlier detection was performed on overall scores, agnostic of condition. For Experiment 1, we recruited 100 participants (49 female, 51 male; age *M* = 27.35 years, *SD* = 5.81), of which 3 were classified as outliers and removed (see Figure S1). The recruitment target was chosen out of convenience, and the resulting sample of 97 participants yields a post-hoc power of 99.8% for a paired contrast (two-tailed) assuming medium effect size (*d* = 0.5), or 49.6% power assuming a weak effect size (*d* = 0.2), based on an *a priori* type I error rate of α = 0.05.

For Experiment 2, we recruited 200 participants (94 female, 106 male; age *M* = 29.85 years, *SD* = 6.16), resulting in 40 participants for each level of the between-subjects factor. This sample size was chosen to ensure at least 80% power (two-tailed) to detect an effect of medium size (*d* ≥ 0.5) within each level, requiring 34 participants per level, plus margin. 6 participants were classified as outliers and removed (see Figure S6). In addition to the *a priori* power considerations regarding the paired comparisons, the power to detect the effect of the between-subjects factor is relevant here. The resulting sample of 194 participants yielded a power of 87.8% to detect a weak correlation (*r* = 0.2). None of the participants in Experiment 2 had also participated in Experiment 1.

In addition to detecting outlying participants, we also screened for outlying experimental stimuli (i.e., scenes), again in a condition-agnostic fashion. There were two outlying items in Experiment 1 and zero outlying items in Experiment 2. These items were discarded from further analysis.

### Data analysis

All analyses were performed using custom-written scripts in Python (Van Rossum & Drake Jr, 1995), using the NumPy (S. van der Walt et al., 2011), SciPy (Virtanen et al., 2020), Pingouin (Vallat, 2018), Pandas (McKinney, 2010), Matplotlib (Hunter, 2007), Seaborn (Michael Waskom et al., 2020), PyMC3 (Salvatier et al., 2016), ArviZ (Kumar et al., 2019), and Bambi (Yarkoni & Westfall, 2016) libraries.

For all pairwise comparisons and correlations, in addition to frequentist statistics like *t*-values, we report Bayes factors quantifying how much more likely the data are under the alternative hypothesis than under the null hypothesis (*BF*_10_). Bayes factors were estimated analytically using noninformative priors: a Cauchy prior on effect size and a Jeffreys prior on variance, using the default Cauchy scale parameter of *r* = 0.707, resulting in a quantity known in the literature as the “JZS Bayes factor” (Jeffreys, 1998; Rouder et al., 2009; Zellner & Siow, 1980). For the three primary paired contrasts of Congruent versus Incongruent, we explored the resulting *t*-based *BF*_10_ values across different plausible levels of Cauchy scale values, to verify that our conclusions do not critically depend on the choice for the default. This control analysis is shown in Figure S12.

In addition to the simple paired comparisons and correlations, we report results from Bayesian hierarchical generalized linear models with full random effects structure. See Supplemental Note 1 for details on the models and sampling scheme. Results from these analyses are primarily summarized using 94% Highest Density Intervals (*HDI*_94_).

Before all statistical analyses (including outlier rejection), we log_10_-transformed bounded variables (reaction time, localization error) to improve normality and stabilize variance.

All tests with *a priori* directional hypotheses report one-tailed *p*- and *BF*_10_-values; tests without *a priori* directional hypotheses are conducted using two-tailed values.

## Results

### Congruency costs in change detection with controlled stimuli

A sample of 100 volunteers participated online in Experiment 1, which was a version of the classical “flicker” change detection paradigm (Rensink et al., 1997) (Figure 1a), with an added localization response. Importantly, scene changes could occur in either a congruent or an incongruent object, where low-level similarities between conditions were matched as much as possible (Öhlschläger & Võ, 2017) (Figure 1b), and stimuli were counterbalanced across participants.

Change detection reaction times were well within the maximum of 7.8 s (*M* = 1,576 ms, *SD* = 234; Figure S1), indicating that participants were able to perform the task successfully. This was further corroborated by the localization error scores, which demonstrate that participants on average were able to locate the item correctly (*M* = 26.8%, *SD* = 9.7, where values < 50% indicates a click inside the key object; Figure S1). Only 2.10% of total responses had a localization error > 100%, and were excluded from all reaction time analyses.

The key hypothesis that this experiment was designed to test is that objects in congruent contexts incur a cost in change detection performance, compared to incongruent contexts. Specifically, as operationalized here, reaction times for congruent trials should be longer than for incongruent trials. This is exactly what we found (*t*_(96)_ = 4.98, *p* < .001, *d* = 0.51, *BF*_10_ = 10722.90; difference *M* = 0.052 log_10_(RT/s), *CI*_95_ = [0.035, 0.070]; *M*_congruent_ = 1.63 s, *M*_incongruent_ = 1.51 s, difference *M* = 0.12 s; Figure 1c). Additionally, we found that localization errors were larger on congruent than on incongruent trials (*t*_(96)_ = 2.44, *p* = .008, *d* = 0.25; difference *M* = 0.020 log_10_, *CI*_95_ = [0.0064, 0.033]; Figure S2), though the evidence for this effect was only moderate (*BF*_10_ = 3.76).

Although the stimulus set we used is highly controlled, stimuli are still instances of natural scenes. This improves the ecological validity of the experiment, but it also means that stimuli are necessarily a sample of all possible scenes that could have been used. This leads to variation in the effect across experimental items (see Figure S3). In addition to the null-hypothesis test (NHT) described above, we therefore conducted a fully Bayesian hierarchical analysis to account for this, in this case analogous to a paired contrast with random intercepts and slopes across both subjects and stimulus items. The results of this analysis corroborate the conclusions of the NHT: reaction times are slower for congruent than for incongruent trials (coefficient posterior *M* = −0.058 log_10_(RT/s), *HDI*_94_ = [−0.10, −0.010]; see Figure S4). Furthermore, the Bayesian analysis allows us to generalize our conclusion further: across both the population from which our participants were drawn, and the population from which our stimuli were drawn, we can be 98.71% certain that the reaction time effect is non-zero. The Bayesian analysis of localization error also yields a corroboration of the analogous NHT, albeit a weaker one (*M* = −0.021 log_10_, *HDI*_94_ = [−0.046, 0.0061]; see Figure S5), with 92.77% probability that the effect is lower than zero.

In summary, Experiment 1 provided strong evidence that change detection is impaired when the changing object is surrounded by semantically congruent, rather than incongruent, contexts. We extend the existing literature by showing that this effect persists even for controlled, matched, stimuli.

### Congruency costs extend to perceptual identification

A key motivation for the present research was to investigate whether congruent surroundings have consequences for the perception of the congruent or incongruent key object itself. We therefore conducted a second experiment, using the same stimuli as in Experiment 1, but with a different, more directly perceptual, task. 200 volunteers participated in Experiment 2, in which they had to identify one of two similar objects as having been present in a previously presented scene (Figure 2a). Context was again included as a factor (Congruent/Incongruent), and we additionally included the factor Probe (Key/Other), resulting in the addition of trials where an incongruent object was present, but was not the target item (Figure 2b).

Overall two-alternative forced choice (2AFC) accuracy was clearly above chance level (*M* = 61.44%, *SD* = 7.72), and reaction times were well within the maximum of 2.5 s (*M* = 1,226 ms; *SD* = 216; Figure S6), indicating that participants were able to perform the task successfully.

We found that 2AFC accuracy was significantly lower for Congruent trials than for Incongruent ones, when focusing on Probe-Key (*t*_(193)_ = −4.49, *p* < .001, *d* = 0.32, *BF*_10_ = 2013.77; difference *M* = −6.18%, *CI*_95_ = [−8.46, −3.90]; Figure 3). There was no difference for Probe-Other (*t*_(193)_ = −0.23, *p* = .591, *d* = 0.02; difference *M* = −0.30%, *CI*_95_ = [−2.45, 1.85]; Figure 3), with the data about six times more likely under the null hypothesis of no difference (*BF*_10_ = 0.16). Reaction time data showed a very similar pattern: responses were slower for Congruent than for Incongruent trials within Probe-Key (*t*_(193)_ = 4.21, *p* < .001, *d* = 0.30, *BF*_10_ = 677.40; difference *M* = 0.053 log_10_(RT/s), *CI*_95_ = [0.032, 0.074]; *M*_congruent_ = 1.15 s, *M*_incongruent_ = 1.09 s, difference *M* = 0.060 s; Figure S7), with no difference in Probe-Other (*t*_(193)_ = −1.03, *p* = .152, *d* = 0.07, *BF*_10_ = 0.27; difference *M* = −0.013 log_10_(RT/s), *CI*_95_ = [−0.035, 0.0080]; *M*_congruent_ = 1.21 s, *M*_incongruent_ = 1.24 s, difference *M* = −0.032 s; Figure S7).

**Figure 3:**
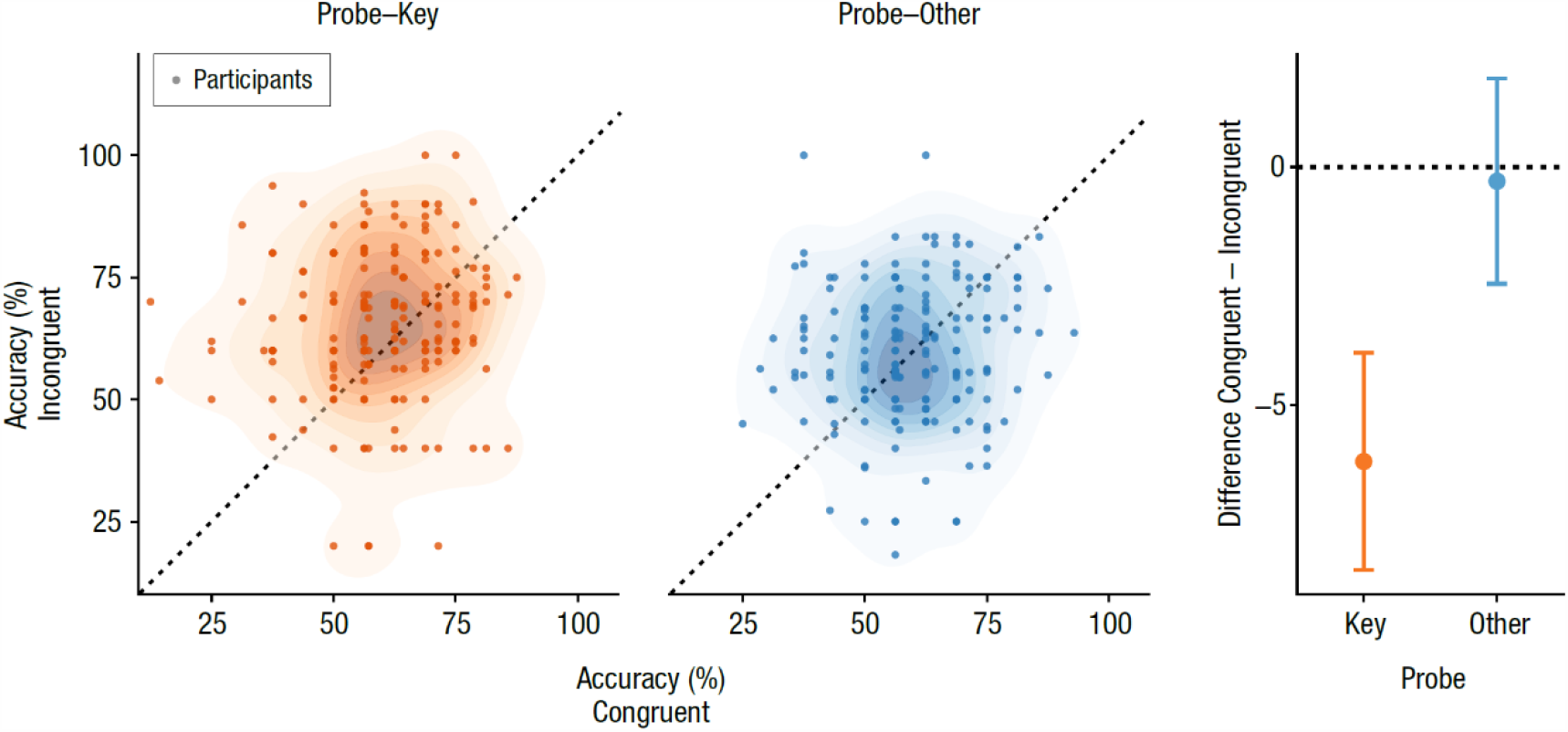
2AFC accuracy results for Experiment 2: responses are more accurate for incongruent scenes. Scatterplots show accuracy in Congruent and Incongruent trials, separately for Probe-Key (orange, left) and Probe-Other (blue, right). Dots are individual participants. Right panel shows mean ± 95% confidence interval of difference scores for both Probe conditions.

Since Experiment 2 constitutes a 2×2 design, it is important to formally test the interaction between the two factors. Additionally, as in Experiment 1, we sought to test the generalization of the observed effects to the population not just of participants, but of experimental items as well (see Figure S8 for effect spread over items). To accomplish these goals, we again conducted a hierarchical Bayesian (logistic) regression analysis of 2AFC accuracy, with the key experimental effect captured by the interaction parameter. We found clear evidence for an interaction effect (*M* = 0.39, *HDI*_94_ = [0.12, 0.65]; see Figure S9), with a 99.66% probability that the parameter exceeds zero, indicating that participants indeed were more accurate for incongruent trials specifically when probed about the (incongruent) key object, and not when probed about another.

The reaction time data showed a very similar pattern: participants were faster on incongruent trials, specifically for Probe-Key (interaction parameter *M* = −0.064 log_10_(RT/s), *HDI*_94_ = [−0.11, −0.014]; see Figure S10), with a probability of 99.24%. Reaction time analysis additionally revealed a main effect of Probe (*M* = −0.060 log_10_(RT/s), *HDI*_94_ = [−0.097, −0.024]), indicating that participants were faster to respond on Probe-Key than on Probe-Other trials (probability 99.88%). It is possible that the probe stimuli in the Other condition were more difficult than those in the Key condition, but given the convincing absence of a main Probe effect in 2AFC accuracy (*HDI*_94_ = [−0.19, 0.30]), we cannot conclude this with certainty.

Taken together, we can conclude with considerable confidence that congruency costs are not limited to perceptually indirect measures like change detection or spatial attention allocation, but extend to object exemplar identification, and thus have genuine perceptual consequences. (See Supplemental Note 2 for an additional result regarding the possible attentional locus of this effect.)

### Congruency costs are not explained by task strategy

In Experiment 1, as in the majority of previous research on object congruence, if an incongruent item was present in a scene, this was always the task-relevant item. Any congruency costs (or: incongruency benefits) might therefore be explained by participants adopting the strategy of always searching for an incongruent object and paying full attention to it. This strategy is only available on incongruent trials, and not on congruent ones, hence potentially causing a general congruency cost. In Experiment 2, we included the Probe-Key/Probe-Other factor specifically to ensure that incongruent items, when present, were not automatically task-relevant (Figure 2b). This factor by itself, in a 2×2 balanced design, already ensures that *p*(*Probe-Key* = 1 | *Incongruent =* 1), or *p*(*Probe-Key* | *Incongruent*) for short, is reduced to 50% (from the typical 100%). For the above analysis, *p*(*Probe-Key* | *Incongruent*) indeed was 50% on the aggregate, yet we observe that congruency costs persist. Therefore, based on the above, we can already conclude that congruency costs are not exclusively observed in settings where *p*(*Probe-Key* | *Incongruent*) = 100%.

It is possible that congruency costs, although not abolished, are still attenuated for lower values of *p*(*Probe-Key* | *Incongruent*). If this were the case, then the strategic concern described above might still be an issue. For Experiment 2, we manipulated *p*(*Probe-Key* | *Incongruent*) across participants to test to what extent the potential confound of task strategy might explain observed congruency costs. We found no effect of *p*(*Probe-Key* | *Incongruent*) on task effects in either 2AFC accuracy (*r*_(192)_ = .11, *p* = .11, *CI*_95_ = [−.028, .25]; Figure 4) or reaction time (*r*_(192)_ = −.027, *p* = .71, *CI*_95_ = [−.16, .11]), with the data about three times more likely under the null hypothesis of no correlation for accuracy (*BF*_10_ = 0.31), and about ten times more likely under the null hypothesis for reaction time (*BF*_10_ = 0.096). Visual inspection (Figure 4) showed that fluctuations in accuracy across *p*(*Probe-Key* | *Incongruent*) were accompanied by opposite fluctuations in reaction time, perhaps suggesting variations in speed/accuracy tradeoff. To account for this, we additionally computed Inverse Efficiency Score (IES) (Vandierendonck, 2017), which also did not show an effect of *p*(*Probe-Key* | *Incongruent*) (*r*_(192)_ = .13, *p* = .057, *CI*_95_ = [−.0042, .27]), though we note that the data here were only about twice as likely under the null hypothesis (*BF*_10_ = 0.54). We finally note that the cell *p*(*Probe-Key* | *Incongruent*) = 17% contains relatively few Incongruent/Probe-Key trials per participant, thus the effect of congruency is estimated less reliably for these participants (also evident in the increased error bars in Figure 4 for this level of the independent variable).

**Figure 4.**
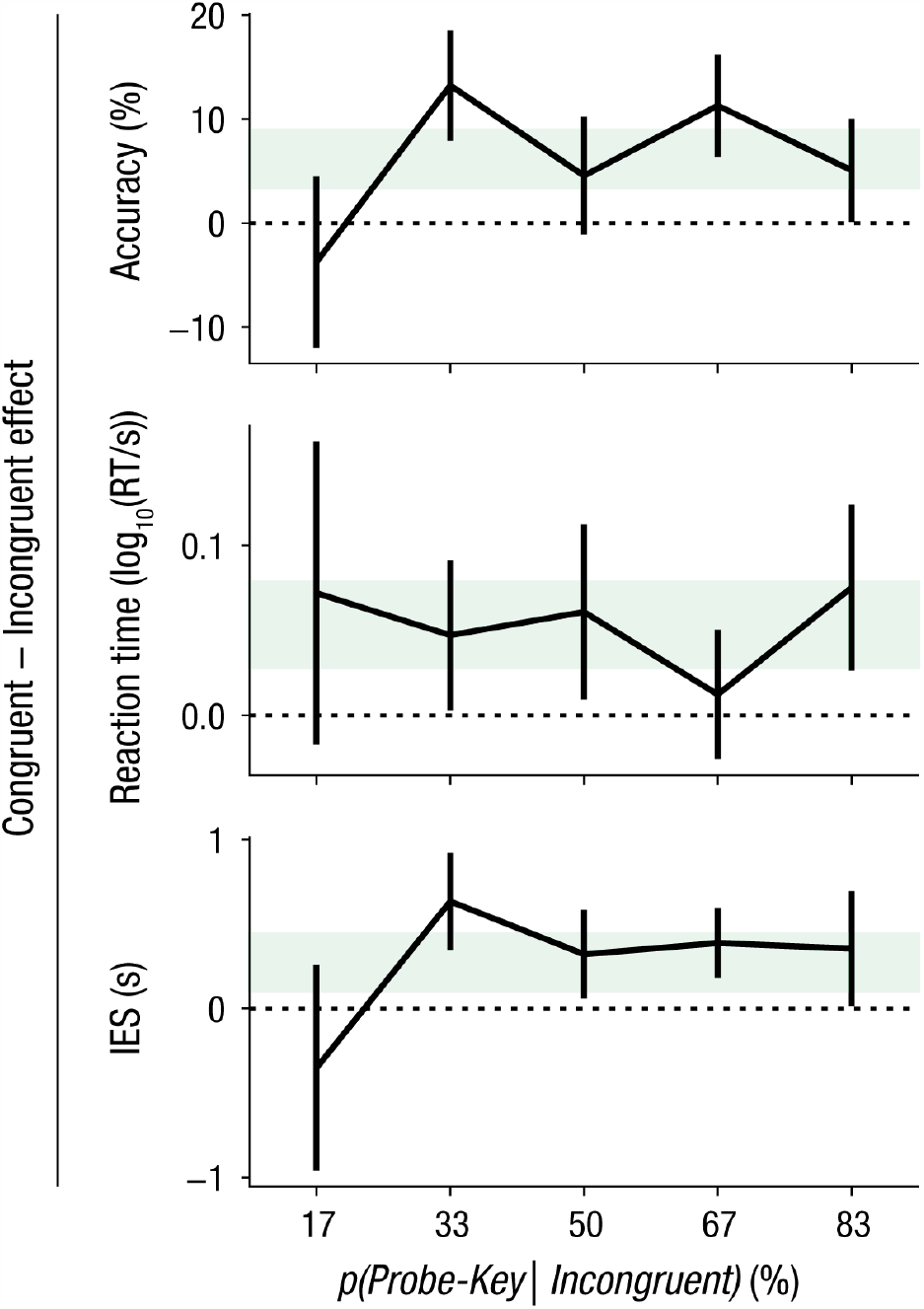
No modulation of congruency effect by relevance manipulation in Experiment 2. Congruent – Incongruent difference scores are shown per level of *p*(*Probe-Key* | *Incongruent*), mean ± 95% confidence interval. Values are shown for the two main dependent variables, 2AFC accuracy and reaction time, as well as for a combined metric, Inverse Efficiency Score (IES). Shading reflects the 95% confidence interval of the effect across the entire sample.

In summary, the persistence of congruency costs in the presence of Probe-Other trials, and in particular the absence of a modulation of congruency costs by increasing task-relevance of incongruent objects, is clear evidence that the congruency cost phenomenon cannot be explained by task-strategic considerations.

### Change detection and exemplar identification tap into overlapping effects

A natural question to ask is whether the congruency cost effects identified in both experiments are related. The two experiments were performed in two different samples of participants, yet used the same stimulus set. Therefore, to answer this question, we examine this relationship across experimental items. We find that, indeed, the experimental effects are related between the two experiments. For the main dependent variables of change detection reaction time (Experiment 1) and 2AFC identification accuracy (Experiment 2), we find clear evidence for a negative correlation across items for congruency difference scores (*r*_(58)_ = −.35, *p* = .003, *CI*_95_ = [−.55, −.10], *BF*_10_ = 11.46; Figure 5; note that the negative direction of the effect is explained by positive (Δ-) accuracy values indicating better performance, while positive (Δ-) reaction time values indicate *worse* performance). Correlations between the other dependent variables corroborated this finding: change detection localization performance was highly correlated with identification reaction time (*r*_(58)_ = .40, *p* < .001, *CI*_95_ = [.16, .59], *BF*_10_ = 43.24), while the other pairwise correlations tended nonsignificantly in the compatible direction (Exp. 1 RT × Exp. 2 RT: *r*_(58)_ = .10, *p* = .21, *CI*_95_ = [−.15, .35], *BF*_10_ = 0.34; Exp. 1 localization error × Exp. 2 2AFC accuracy *r*_(58)_ = −.21, *p* = .056, *CI*_95_ = [−.44, .05], *BF*_10_ = 1.05; Figure 5).

**Figure 5.**
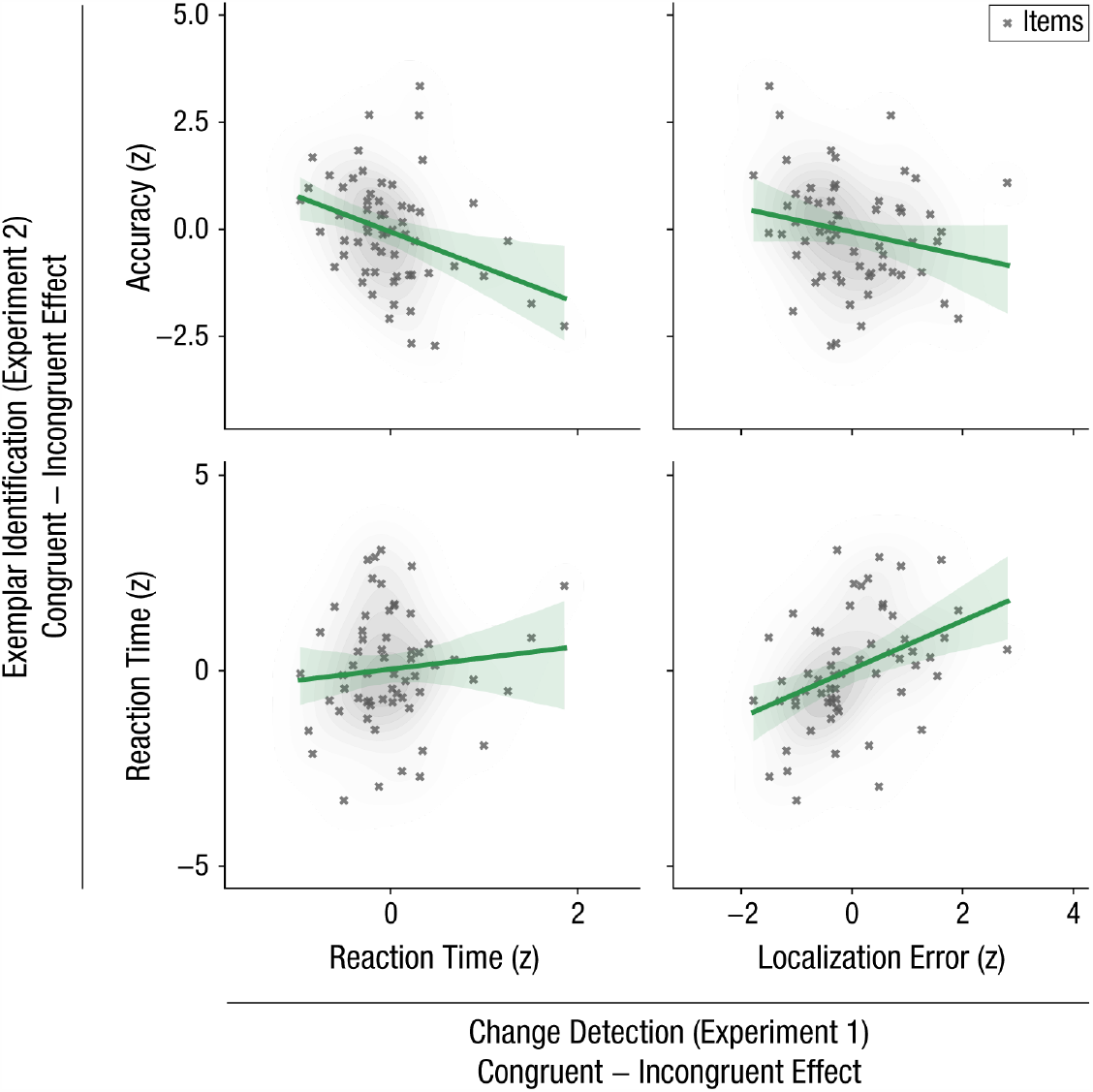
Contextual costs are related between the two experiments. Change detection reaction time and localization error (x-axes) versus exemplar 2AFC identification accuracy and reaction time (y-axes) for the different items. Lines indicate best-fitting regression line of 2AFC effect onto change detection effect, shading indicates 95% confidence interval of regression line.

We can thus conclude that the two perceptual tasks, change detection in Experiment 1, and exemplar identification in Experiment 2, tap into overlapping effects.

### Subjective congruency ratings partly explain behavioural performance

Within the incongruent scenes, there is variation in the extent to which an object might be considered incongruent. This variation has previously been established by the authors of the original publication on the stimulus set we used: a separate sample of observers independently rated each of the scene stimuli (Öhlschläger & Võ, 2017). As a final, exploratory, analysis, we asked whether this variation in subjective (in)congruency ratings might explain (part of) the congruency effects we observed. Specifically, we looked at the correlation between subjective inconsistency ratings (where higher means more *in*consistent) and behavioural performance across the incongruent scenes. Here, as in the section above, we leveraged variation across experimental items, rather than across participants.

Change detection reaction times in Experiment 1 were not correlated with inconsistency rating (*r*_(58)_ = .05, *p* = .358, *CI*_95_ = [−.21, .30], *BF*_10_ = 0.12; Figure 6). However, exemplar identification 2AFC accuracy scores in Experiment 2 were significantly correlated with inconsistency ratings, with higher subjective inconsistency corresponding to better performance (*r*_(60)_ = .32, *p* = .005, *CI*_95_ = [.08, .53], *BF*_10_ = 7.67; Figure 6). Neither of the secondary dependent variables in the two experiments was significantly correlated with inconsistency ratings (both |*r*| < .15; p > .1; *BF*_10_ < 0.2; Figure S11).

**Figure 6.**
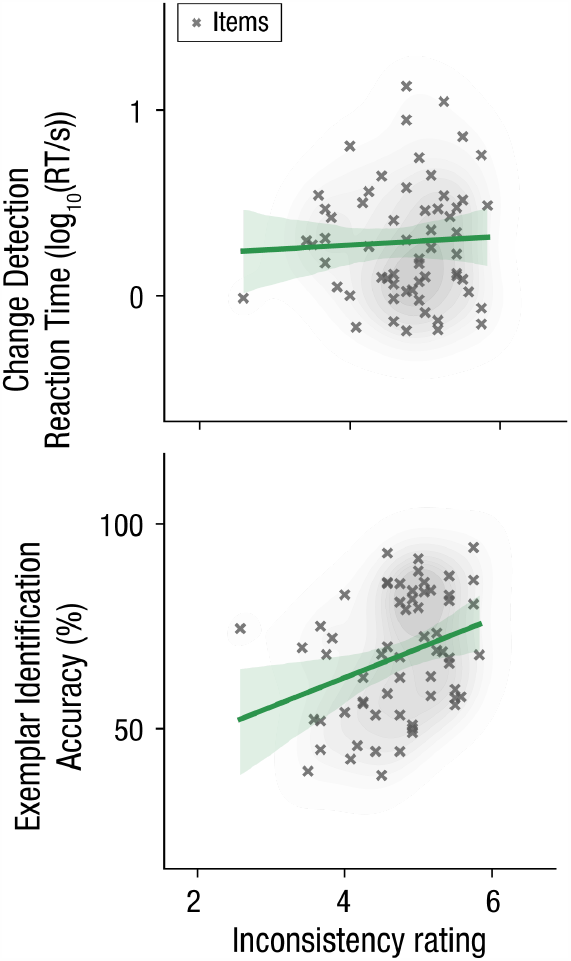
Subjective inconsistency ratings partly explain behavioural performance. Subjective inconsistency ratings for the different experimental items, provided by an independent sample of observers, as collected by Öhlschäger & Võ (2017). Ratings are shown on the x-axis, with higher scores indicating stronger inconsistency (i.e., lower object/scene consistency). Although these ratings appear unrelated to change detection performance in Experiment 1, they are significantly correlated with 2AFC identification performance in Experiment 2. Crosses are individual items (incongruent scenes only). Lines indicate best-fitting regression, shading indicates 95% confidence interval of regression line.

We can conclude that objects within scenes that are rated as subjectively more incongruent by independent observers are easier to identify (Experiment 2), but changes in those objects are not detected faster (Experiment 1).

## Discussion

Using two large-sample behavioural experiments, we tested a counterintuitive consequence of the theory of predictive coding: that scene-induced priors can lead to impaired perception of scene-congruent objects. In Experiment 1, we replicated the established congruency cost effect from the change detection literature, while crucially controlling for potential stimulus confounds previously uncontrolled for. More importantly, in Experiment 2, using a task requiring object exemplar identification, we demonstrated that congruent surroundings impair perception of key objects themselves. By manipulating the proportion of incongruent objects that were relevant, we were able to show that congruency costs are not due to participant strategy, but rather reflect an automatic perceptual phenomenon.

Previous studies have reported slower change detection responses for changes in congruent objects than for changes in incongruent ones (Hollingworth & Henderson, 2000; LaPointe et al., 2013; Mack et al., 2017). Eye tracking studies have additionally reported that scene-incongruent objects are fixated earlier than congruent ones (Bonitz & Gordon, 2008; Loftus & Mackworth, 1978; Underwood et al., 2007). It has been debated to what extent these effects are truly due to *semantic* congruence, or might instead be better explained by low-level visual features (e.g., local contrast) differing between conditions (Underwood & Foulsham, 2006; Võ & Henderson, 2009). In the present study, we used a stimulus database designed to be highly balanced between the semantically (in)congruent conditions (Öhlschläger & Võ, 2017). A further advantage of these stimuli is that they were all actual photographs, with no reliance on digital image editing techniques to ‘transplant’ objects from one context to another. Such digital editing, even when used carefully, might introduce local inconsistencies (e.g. in lighting) that are not exclusively semantic. The results for Experiment 1 demonstrate that congruency costs are evident in change detection even for such controlled stimuli, ruling out worries that this semantic effect might not truly be semantic at all.

An additional potential concern is that of task strategy. In a typical experimental design with congruent and incongruent trials, if an incongruent object is present in a scene, then this is very likely task-relevant (e.g., the locus of change in change detection, or specifically memorable in a memory task). Participants might therefore decide to always look for an incongruent object, as this will be beneficial in half the trials (and thus in general). This strategy is only available on incongruent trials, thereby leading to a behavioural benefit in that condition. We quantified this potential strategic effect in Experiment 2, by including trials on which an incongruent object was present, but not task-relevant. Importantly, we manipulated the proportion of trials on which a presented incongruent object was relevant across participants, and found no modulation of congruency costs by this proportion. We can therefore conclude that these congruency costs arise automatically, independent from (deliberate or unconscious) strategic choices. This conclusion is in line with Hollingworth & Henderson’s (2000) work, which included a similar manipulation to our Probe-Key/Other factor, and crucially extends it with the manipulation of relevance probability.

As described, previous work has identified congruency costs (or, equivalently: incongruency benefits) before in perceptually indirect, attentional measures. Perhaps the most important advance made by Experiment 2 over earlier work is that it probed the consequences of scene congruency for object perception itself, through a different task than previously used, namely exemplar identification. We used accuracy in a bias-free discrimination task as dependent variable, thereby enabling us to test whether scene congruency affects perceptual encoding, independently of decisional and response biases. Also here we found clear evidence for congruency costs, which was furthermore related to the effect observed during change detection. Congruency costs thus not only reflect non-specific biases or some peculiarity of change detection, but indeed appear generally during the process of perceiving objects within scenes.

It is possible that the effects we observed in Experiment 2 were, at least partly, mediated by observers directing their attention (overtly or covertly) to incongruent objects more frequently than to congruent ones. This would be in line with some interpretations of predictive coding theory, upon which prediction errors elicited by incongruent objects increase the salience of these objects and thus attract attention ((Den Ouden et al., 2012; Feldman & Friston, 2010); see also the next paragraphs). Since we did not measure eye movements, our results do not speak directly for or against this interpretation. The absence of a congruency effect when the key object was irrelevant (Probe-Other trials; see also Supplemental Note 2) provides some circumstantial evidence against spatial attention being the only factor at play here, but further research is needed to understand whether congruency costs, as observed in Experiment 2, are the cause or the consequence of attentional orienting.

It has been claimed that attentional orienting is best understood as reflecting ‘active sampling’ within the framework of predictive coding (Dey & Gottlieb, 2019; Feldman & Friston, 2010). An implication of the predictive coding scheme is that newly incoming data that are unlikely under some prior expectation elicit larger prediction error responses than those stimuli for which prior probability is high. Resolving these errors entails closer inspection (increased attentional sampling) of the corresponding stimuli. An important source of prior information in natural visual perception is the gist of a scene (Bar, 2004). Congruent objects are easily accommodated by such a scene-induced prior, whereas incongruent objects should be inspected more closely before a coherent posterior is possible. This difference in amount of processing should have consequences for subjective awareness (Anzulewicz et al., 2015; Windey et al., 2013). The main motivation for the present study was to test the key hypothesis that gist-induced priors lead to impaired perception of conguent items. Our results corroborate this hypothesis.

The Bayesian brain hypothesis is often invoked to explain congruency *benefits* in perception (de Lange et al., 2018), for which ample empirical evidence exists. Nevertheless, we report robust congruency *costs*. These seemingly contradictory claims are reconciled by noting that congruency benefits in scene perception are typically reported for tasks where the gist-induced prior is, by itself, already helpful for behaviour. Once the gist of a scene is recognized as corresponding to a barbershop, observers will more likely identify objects inside that scene as hairdryers, even independently from the associated sensory input. Similarly, observers can use gist-based knowledge to efficiently guide a search for a hairdryer; knowledge that is unavailable when searching for a hammer in the same scene. In contrast to category identification or visual search, change detection is a non-semantic task, where scene gist cannot inform the judgement to be made. We deliberately designed Experiment 2 to similarly involve a judgement orthogonal to any scene-induced prior, and, as predicted, this revealed congruency costs, rather than benefits.

The apparent paradox between congruency benefits in some, and congruency costs in other settings (or, more generally: predictive ‘upweighting’ versus ‘cancellation’) is an active area of debate and research. A recently proposed ‘opposing process’ model aims to resolve this paradox by asserting that perception is initially biased towards a priori likely stimuli, while later processes shift this balance towards (high-precision) unexpected (and thus informative) inputs (Press et al., 2020). Our observations can be neatly accommodated within this framework: high-precision sensory input corresponding to incongruent objects results in informative prediction errors, thus ‘divert[ing] extra perceptual processing resources to the unexpected’ (Press et al., 2020, p. 18), leading to the observed impaired perception of congruent objects.

We finally would like to emphasize that, in addition to presenting frequentist null hypothesis tests, we consistently tested our key hypotheses using Bayesian hierarchical models with full random effects. This allows us to formally generalize our conclusions not just to the population from which our participants were drawn, but also to the population from which our *stimuli* were drawn (Arnqvist, 2020; Yarkoni, 2019), lending further support to the generality of our findings.

In summary, we tested an important and seemingly counterintuitive hypothesis of the influential theory of predictive coding: that prior information due to real-world scene gist leads to reduced processing of objects congruent within such scenes. Across two experiments, with distinct experimental tasks, we found clear evidence in favour of this hypothesis, thereby furthering our understanding of how prior knowledge interacts with sensory input to yield real-world percepts and guide behaviour.

### Open Practices Statement

All experimental stimuli and collected data for both experiments are fully open and can be freely downloaded from the [INSTITUTE] Repository [url available at publication]. All analysis code is released under the GNU General Public License v3.0 and available from GitHub [url available at publication]. Neither of the experiments described in this article was formally preregistered.

## Supplemental Information

### Supplemental Note 1 – Bayesian data analysis

We constructed Bayesian hierarchical generalized linear models (GLMs) for both experiments and for both DVs per experiment. For log-reaction time (log-RT) and log-localization error, we used a Gaussian family and identity link function, whereas for accuracy, we used a Bernoulli family and logit link. All experimental factors of interest included the full random effects structure (both intercepts and slopes) for subjects as well as for items. For Experiment 1, this yields the following GLM equation:

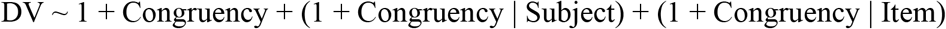

For Experiment 2, this yields the following:

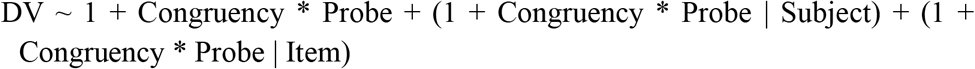

Models were constructed using Bambi, which was also responsible for setting appropriate weakly informative priors. Posterior probability distributions were obtained using Markov Chain Monte Carlo (MCMC) based on the No U-Turn Sampler (NUTS), as implemented in PyMC3. Four chains were sampled for each model, with 3,000 samples per chain, after a 6,000-sample tuning period, using a target acceptance ratio of 90%. Starting values were determined using Automatic Differentiation Variational Inference (ADVI; option init = ‘advi+adapt_diag’), run for 35,000 time steps or until plateau. We checked chain convergence through visual inspection, as well as through the Gelman-Rubin statistic 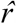. Posterior results are primarily summarized using 94% Highest Density Intervals (*HDI*_94_), based on the combined samples of all chains. Probability of parameters lying above/below a critical value is summarized by the proportion of combined samples above/below that value.

### Supplemental Note 2 – Congruency costs might not always depend on spatial attention

In Experiment 2, if the congruency cost in the Probe-Key condition is due to an incongruent item ‘grabbing attention’ (either through foveation or covertly) more strongly than a congruent item, then we would expect a congruency *benefit* in the Probe-Other condition. Also in that condition, a by definition irrelevant incongruent item (present only in the incongruent condition) would grab attention, thereby impairing performance on discriminating the (always congruent) Probe-Other target item in the scene. As described in Results, our data suggest that the existence of such an effect in Probe-Other trials is unlikely.

As a further test of the attentional locus of this effect, we might ask whether stimulus items that yield a strong Probe-Key congruency effect, also yield a strong Probe-Other congruency effect. Such a correlation might exist even in the absence of a Probe-Other congruency effect in the average. However, also here we find evidence to the contrary, both for 2AFC accuracy (*r*_(60)_ = −.094, *p* = .235, *CI*_95_ = [−0.34, 0.16], *BF*_10_ = 0.31) and for reaction times (*r*_(60)_ = .13, *p* = .154, *CI*_95_ = [−0.12, 0.37], *BF*_10_ = 0.083).

Taken together, the absence of a Congruency effect in Probe-Other, as well as the absence of a relationship between Probe-Key and Probe-Other effects, raise the interesting possibility that identified congruency costs in the exemplar identification task are not entirely mediated by attentional factors. However, given the large body of literature interpreting congruency costs as attentional in other tasks (see Introduction), we do not wish to make strong conclusions here; particularly since we did not record eye movements.

---

**Figure S1.**
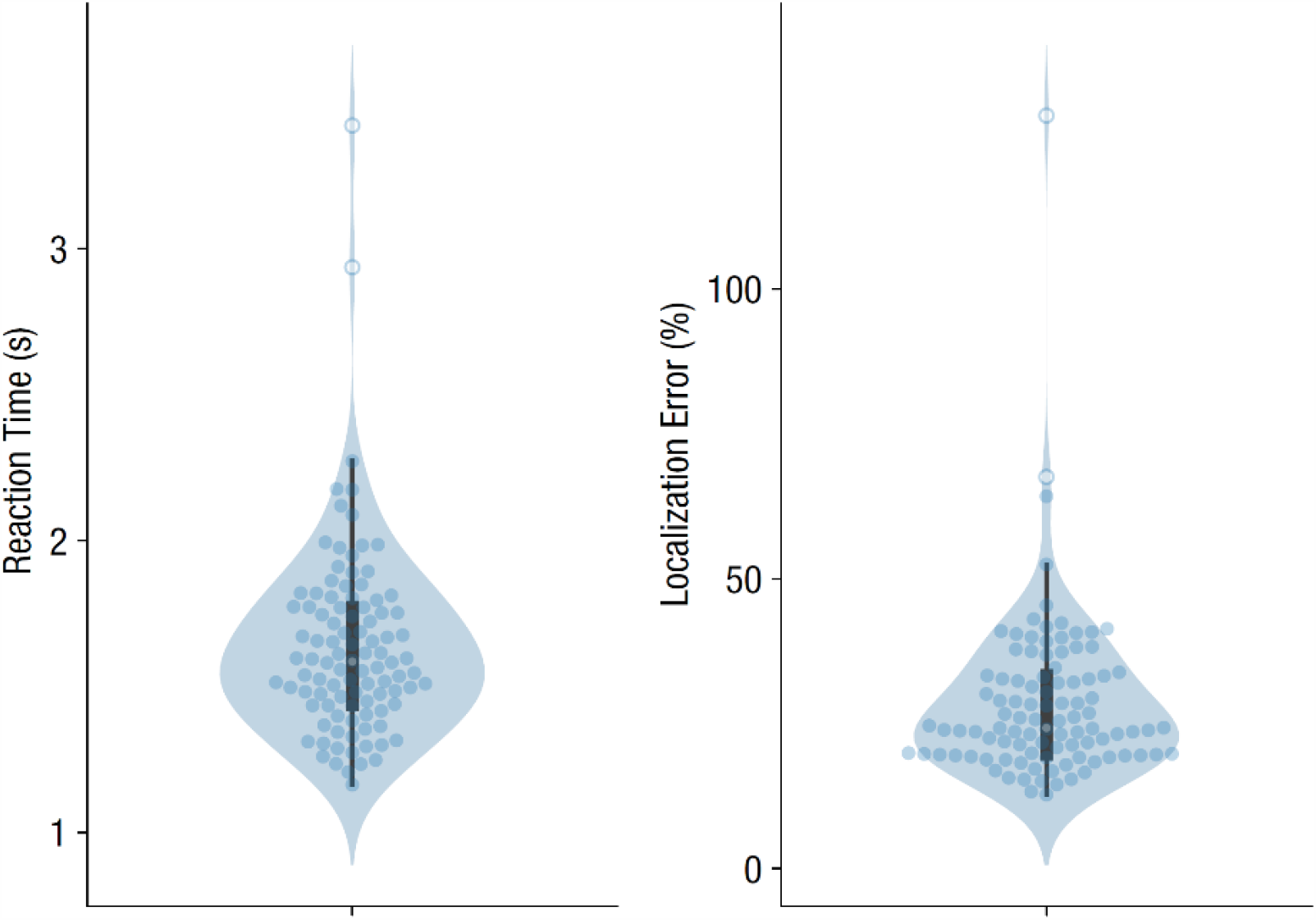
Overall reaction time and localization error distributions for Experiment 1. Dots are participants; hollow circles are outliers (removed from all analyses).

**Figure S2.**
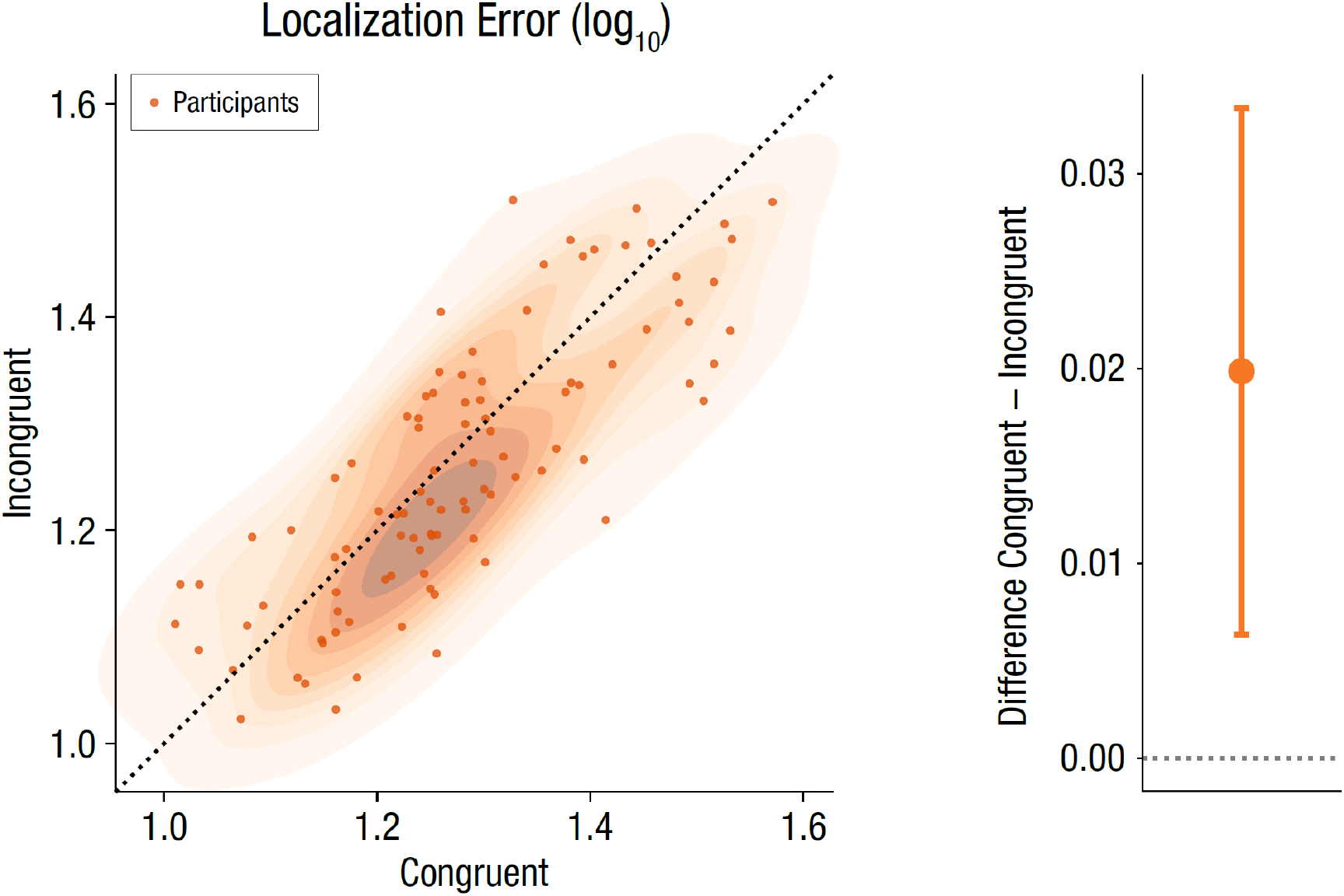
Localization errors across all participants in Experiment 1, for Congruent and Incongruent trials (analogous to Figure 1c). Full distribution in scatterplot, mean ± 95% confidence interval of difference scores on the right.

**Figure S3.**
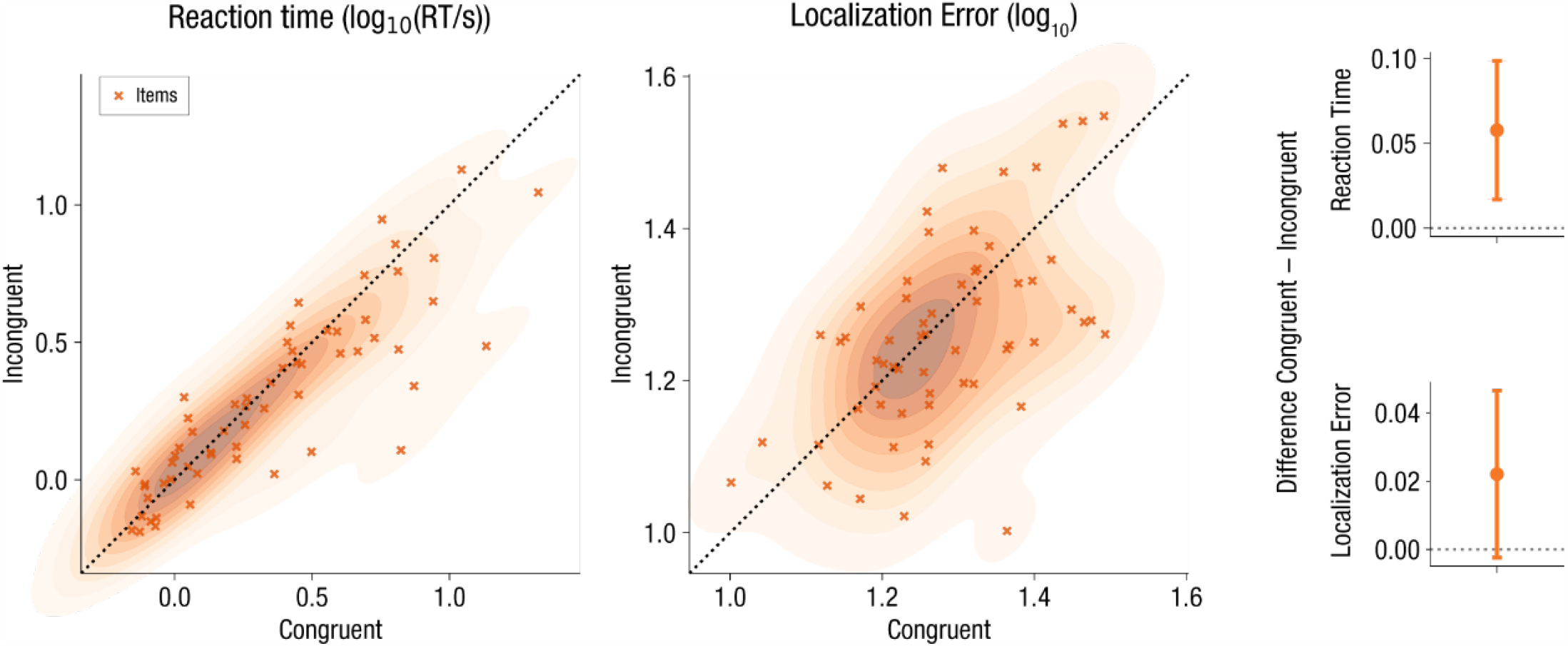
Reaction times and localization errors across experimental items (averaged over participants) for Experiment 1. Crosses are individual items. Full distributions in scatterplot, mean ± 95% confidence interval of difference scores on the right.

**Figure S4.**
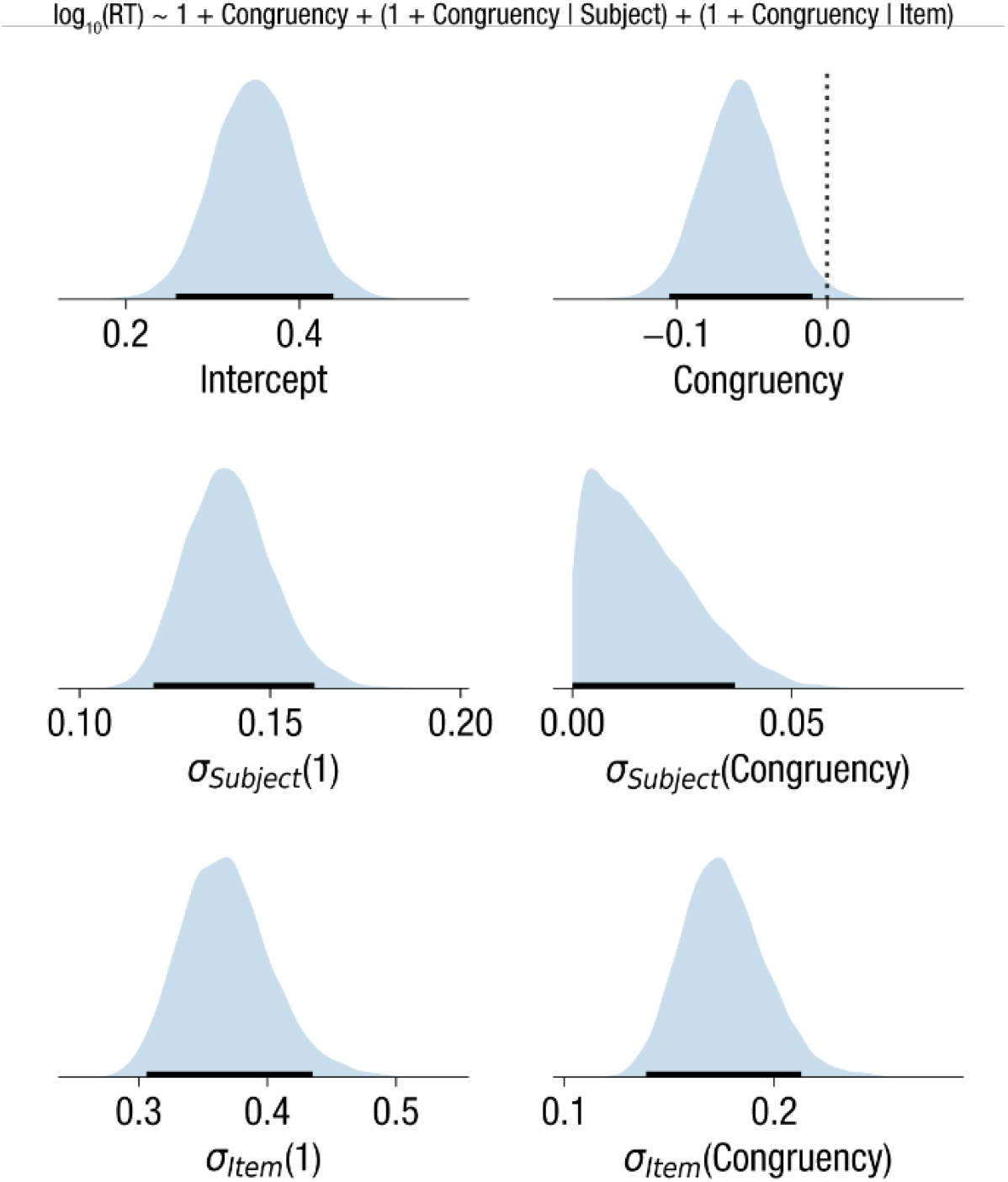
Posterior distribution after MCMC sampling of Bayesian regression model for reaction time in Experiment 1. Top row shows parameter estimates for fixed effects coefficients, middle row corresponds to subject-level random effects (i.e. standard deviation of effect over subjects), bottom row corresponds to item-level random effects. Intercepts and slopes for individual items/subjects are not shown (as these are very many), only the parameters related to their spread are included. Shading reflects a kernel density estimate of the marginal full posterior for one parameter; black horizontal bars indicate 94% HDI; dashed vertical lines, if present, correspond to a reference value (i.e., 0 for a fixed effect).

**Figure S5.**
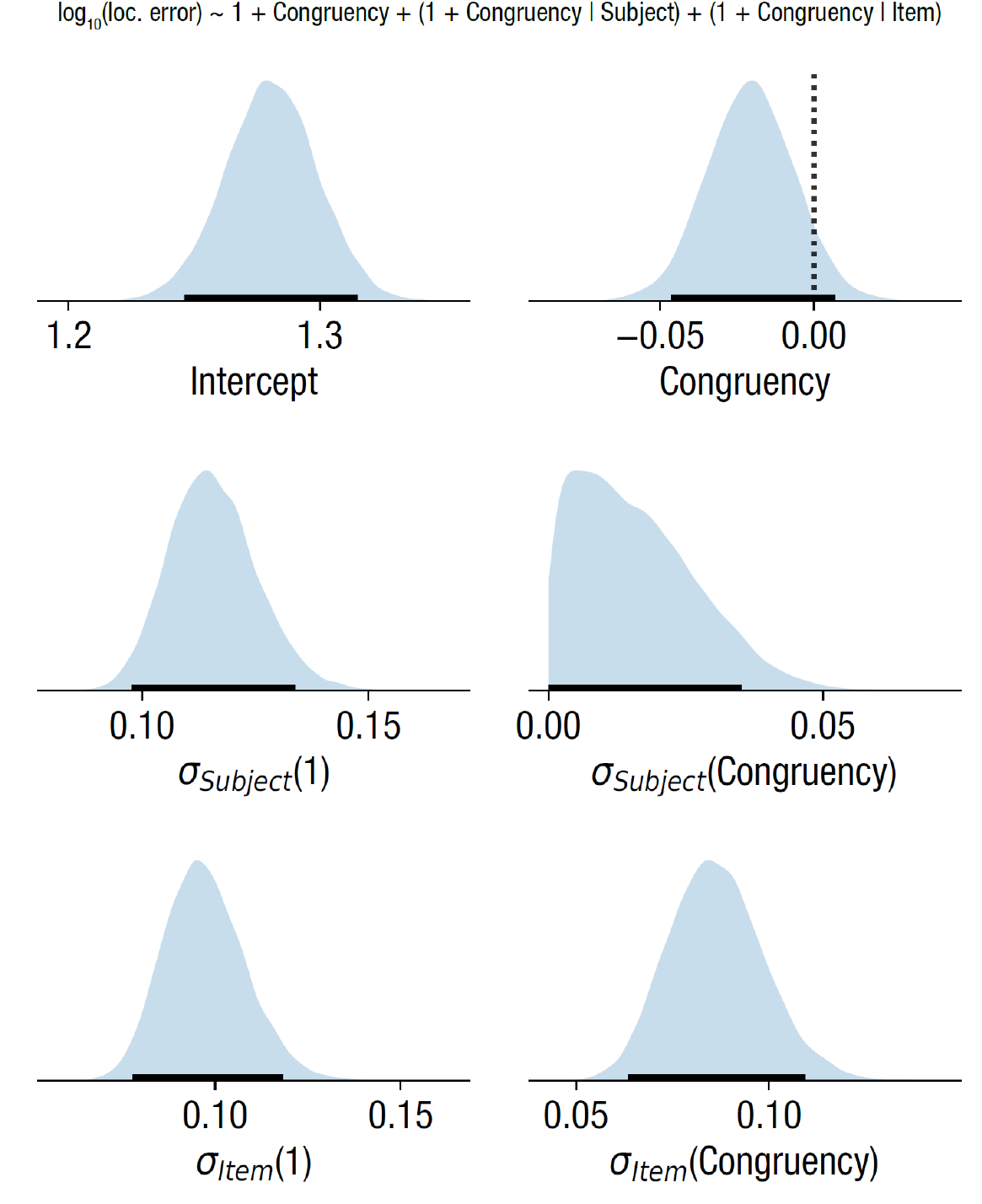
Posterior distribution after MCMC sampling of Bayesian regression model for localization error in Experiment 1. All panels and conventions as in Figure S4.

**Figure S6.**
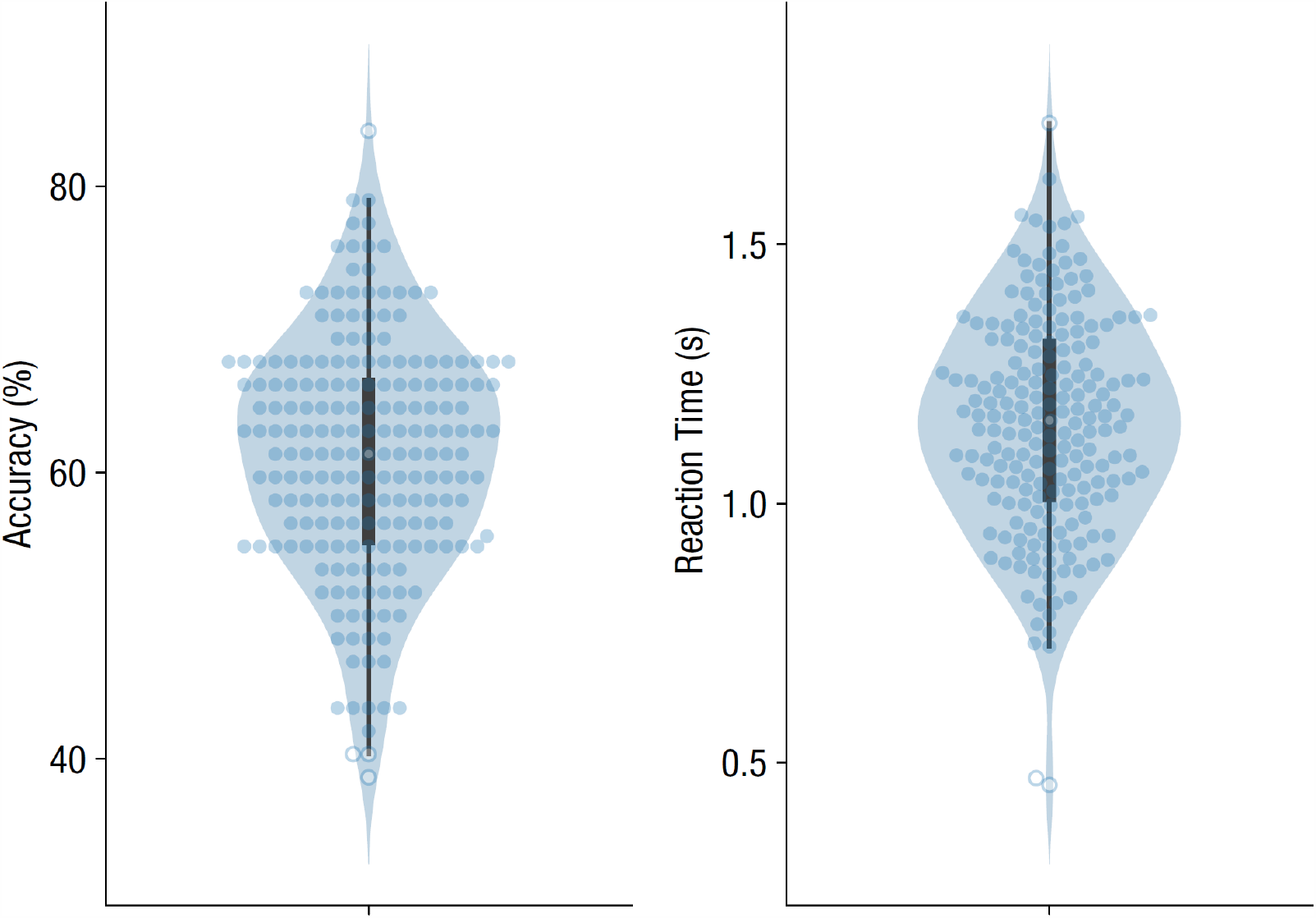
Overall accuracy and reaction time distributions for Experiment 2. Dots are participants; hollow circles are outliers (removed from all analyses).

**Figure S7.**
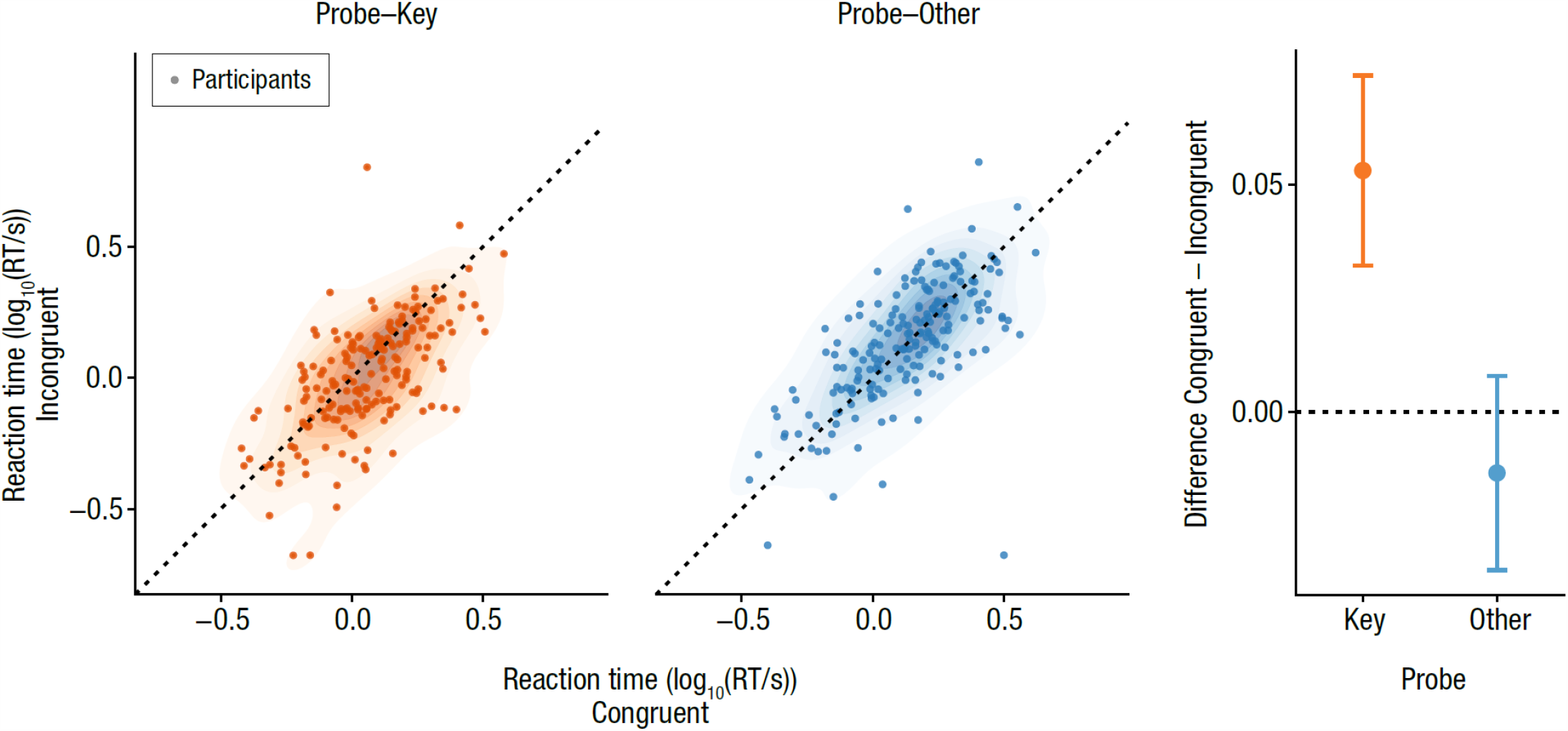
2AFC reaction times in Congruent and Incongruent trials, separately for Probe-Key (orange, left) and Probe-Other (blue, right). Dots are individual participants. Right panel shows mean ± 95% confidence interval of difference scores for both Probe conditions. (Presentation analogous to Figure 3.)

**Figure S8.**
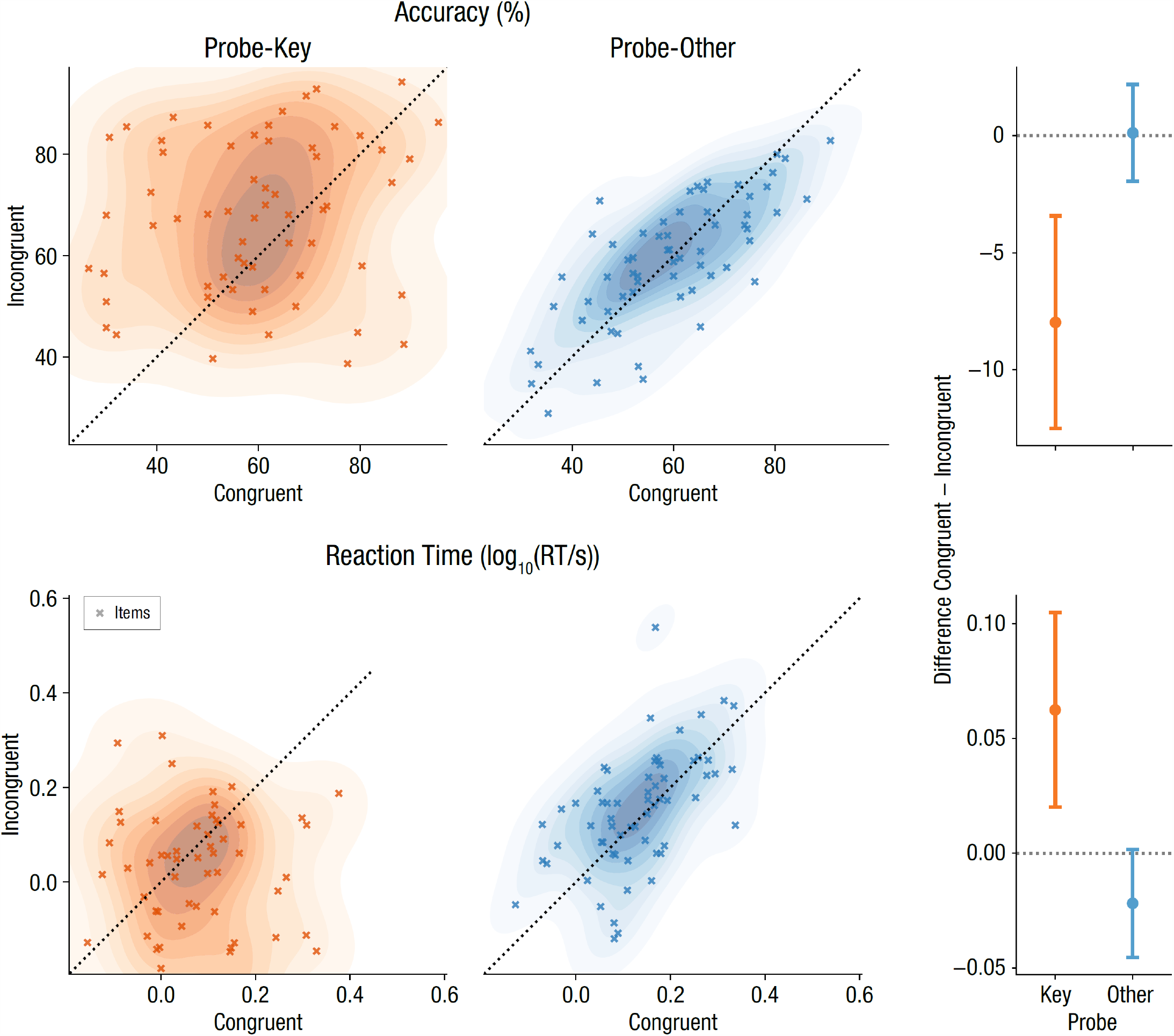
2AFC accuracy (top) and reaction times (bottom) for Experiment 2, across experimental items (averaged over participants), for the Incongruent/Congruent and Probe-Key/Probe-Other conditions. Crosses are individual items. Full distributions in scatterplots, mean ± 95% confidence interval of difference scores in right panels.

**Figure S9.**
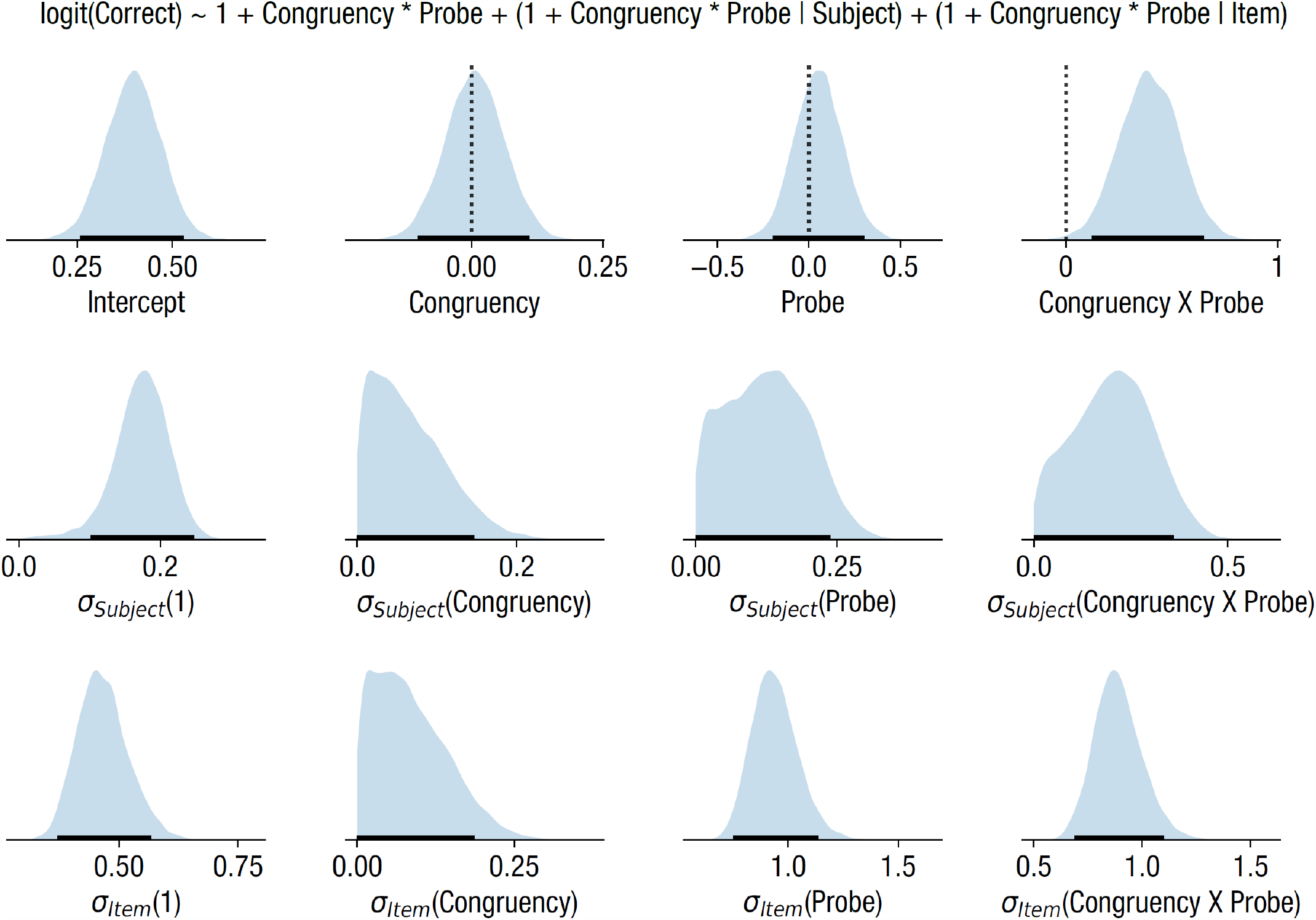
Posterior distribution after MCMC sampling of Bayesian logistic regression model for accuracy in Experiment 2. All panels and conventions as in Figure S4.

**Figure S10.**
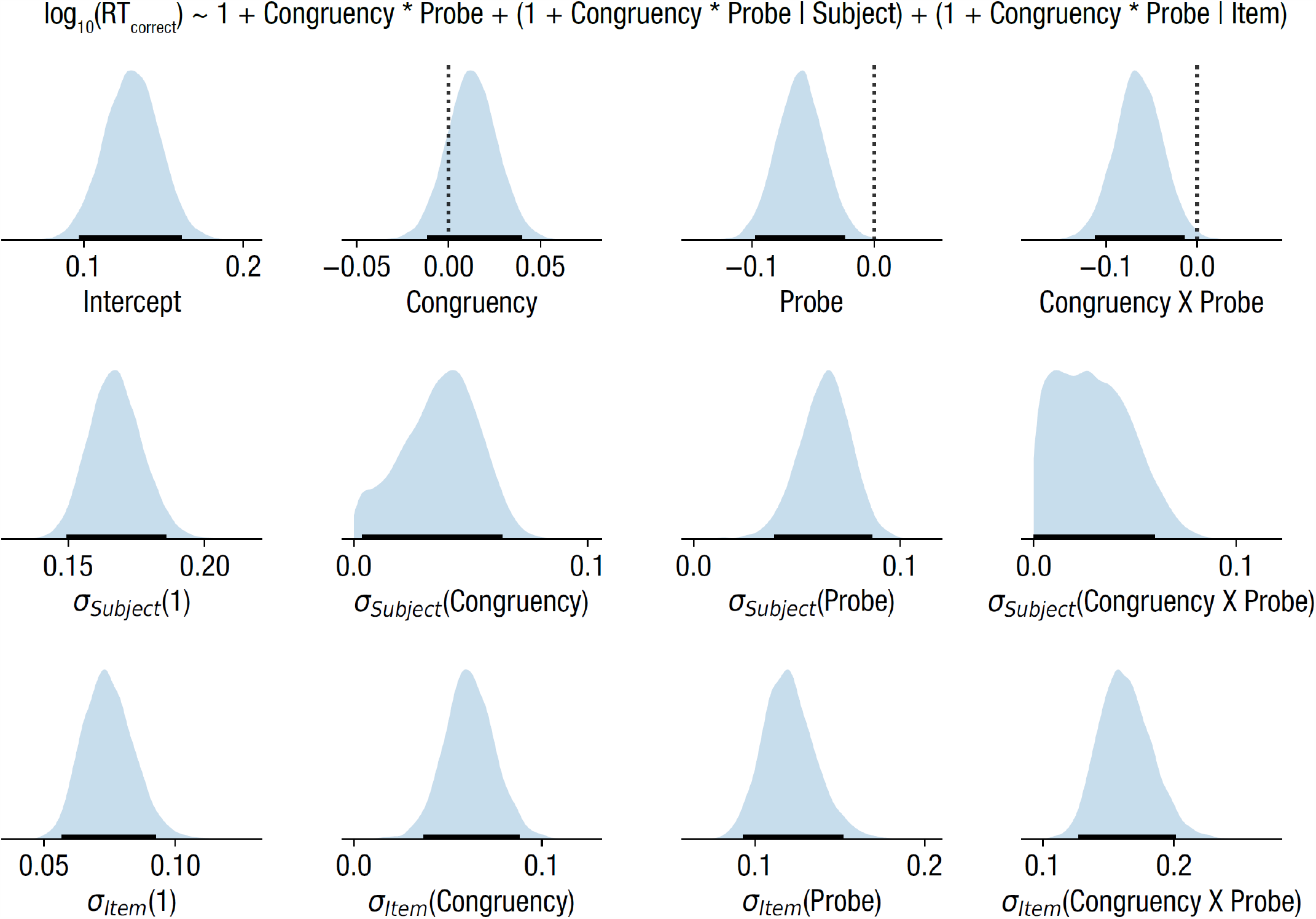
Posterior distribution after MCMC sampling of Bayesian regression model for reaction times in Experiment 2. All panels and conventions as in Figure S4.

**Figure S11.**
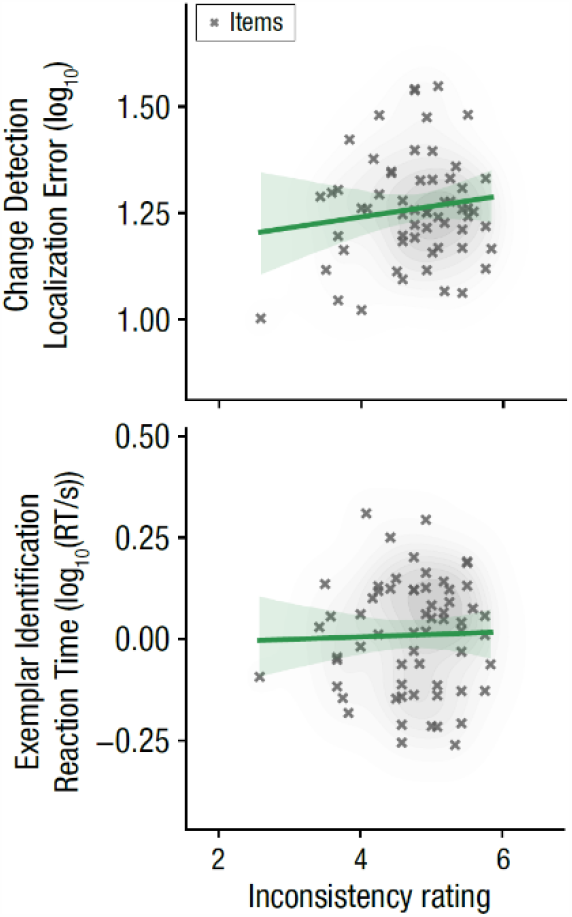
Correlations of secondary dependent variables with subjective inconsistency ratings across all incongruent items. Lines indicate best-fitting regression, shading indicates 95% confidence interval of regression line.

**Figure S12.**
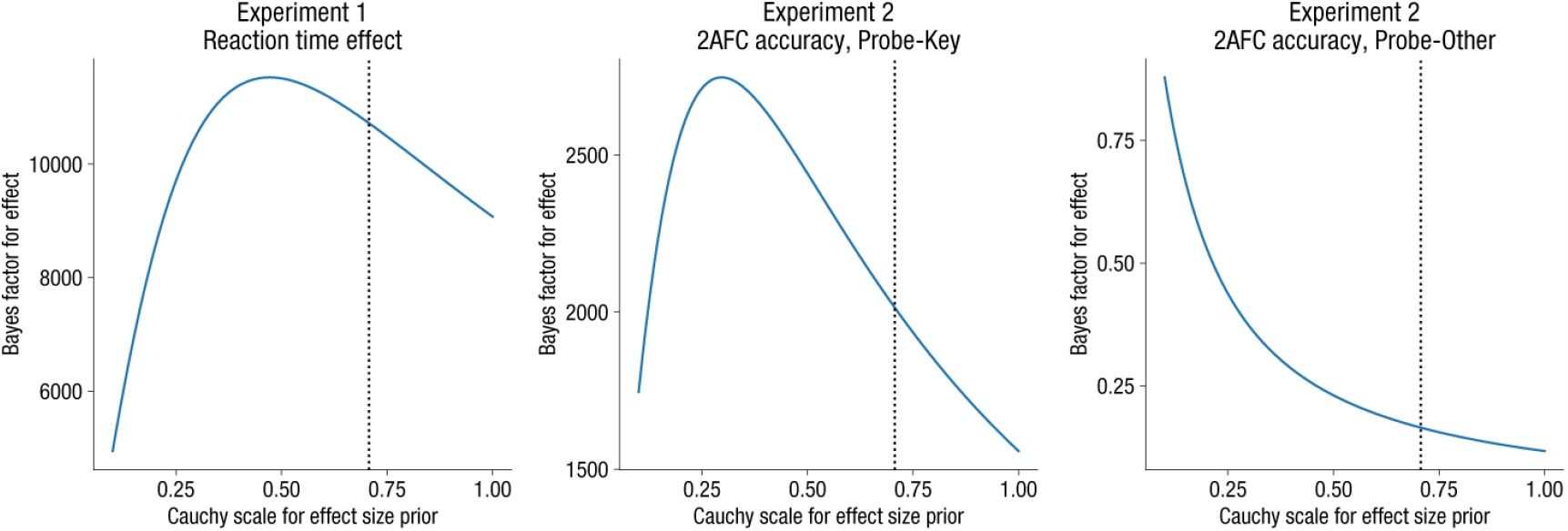
Robustness analysis for *t*-statistic-based Bayes factors (*BF*_10_). *BF*_10_ is shown as a function of the scale parameter *r* used for the Cauchy prior over effect sizes. Different panels correspond to the three primary paired hypotheses tested in the manuscript (see panel headings). The dashed vertical line corresponds to the default scale parameter *r* = 0.707 used for all reported analyses.

## Notes

### Competing Interest Statement

The authors have declared no competing interest.

